# Plastid anionic lipids are required for membrane development and protochlorophyllide synthesis in etioplasts

**DOI:** 10.1101/2023.05.16.541020

**Authors:** Akiko Yoshihara, Keiko Kobayashi, Noriko Nagata, Sho Fujii, Hajime Wada, Koichi Kobayashi

**Author notes:** (Corresponding Author) **Corresponding author:** Faculty of Liberal Arts, Science and Global Education, Osaka Metropolitan University, 1-1 Gakuen-cho, Naka-ku, Sakai, Osaka 599-8531, Japan.

## Abstract

Dark-germinated angiosperms develop the chloroplast precursors called etioplasts in cotyledon cells. Etioplasts develop lattice membrane structures called prolamellar bodies (PLBs), where the chlorophyll intermediate protochlorophyllide (Pchlide) forms a ternary complex with NADPH and light-dependent NADPH-Pchlide oxidoreductase (LPOR). The lipid bilayers of etioplast membranes are mainly composed of galactolipids, which play important roles in membrane-associated processes in etioplasts. Although etioplast membranes also contain two anionic lipids, phosphatidylglycerol (PG) and sulfoquinovosyldiacylglycerol (SQDG), the roles of these anionic lipids are unknown. To reveal the importance of PG and SQDG for the development of etioplasts, we characterized etiolated Arabidopsis mutants deficient in the biosynthesis of PG and SQDG. A partial deficiency in PG biosynthesis loosened the lattice structure of PLBs and impaired the insertion of Mg^2+^ into protoporphyrin IX, leading to a significant decrease in Pchlide content. Although a complete lack of SQDG biosynthesis did not notably affect both PLB formation and Pchlide biosynthesis, the lack of SQDG in addition to the partial deficiency of PG caused strong impairments of these processes. The results suggested that PG is required for PLB formation and Pchlide biosynthesis, whereas SQDG plays an auxiliary role in these processes. Notably, the PG deficiency and the lack of SQDG oppositely affected the dynamics of LPOR complexes after photoconversion, suggesting different involvements of PG and SQDG in the organization of LPOR complexes. Our data demonstrate pleiotropic roles of anionic lipids in etioplast development.

## INTRODUCTION

In plants, chlorophyll (Chl) is synthesized within plastids in coordination with the development of photosynthetic machinery (Wang and Grimm, 2021). Chl biosynthesis begins with the formation of 5-aminolevulinic acid (ALA) from glutamyl-tRNA^Gul^, which is the rate-limiting step for the regulation of the entire pathway (Tanaka et al., 2011). ALA is then converted to protoporphyrin IX (Proto IX), the last common intermediate of heme and Chl biosynthesis, through multiple enzymatic steps. For Chl biosynthesis, Mg^2+^ is inserted into Proto IX by Mg-chelatase to yield Mg-Proto IX, which is further converted to Mg-Proto IX monomethylester (Mg-Proto IX ME) by S-adenosyl-L-Met:Mg-Proto IX methyltransferase (MgMT) and to protochlorophyllide (Pchlide) by Mg-Proto IX ME cyclase (MgCY). Pchlide is then reduced and phytylated, resulting in Chls.

In cyanobacteria and plants, there are two types of NADPH-Pchlide oxidoreductase (POR) for converting Pchlide to chlorophyllide (Chlide), the immediate precursor of Chl (Masuda, 2008). Whereas dark-operative POR can reduce Pchlide to Chlide in the absence of light, light-dependent POR (LPOR) absolutely requires light for its reaction. Angiosperms have only LPOR, so they accumulate Pchlide in plastids called etioplasts when germinated in the dark (Solymosi and Schoefs, 2010). Etioplasts are chloroplast precursors containing 3D lattice membrane structures called prolamellar bodies (PLBs) and flattened lamellar prothylakoids (PTs). Pchlide in etioplasts forms ternary complexes with LPOR and NADPH, most of which further oligomerize and cover the membrane tubules of PLBs in a helical fashion (Floris and Kühlbrandt, 2021; Nguyen et al., 2021). The Pchlide bound by LPOR at the active site (photoactive Pchlide) is immediately converted to Chlide with flash illumination, while the rest of the Pchlide (nonphotoactive Pchlide) remains unchanged (Schoefs and Franck, 2003). The nonphotoactive and photoactive Pchlides can be distinguished by their different fluorescence spectra at 77K with emission peaks at around 633 and 655 nm, respectively (Schoefs, 2001). The photoconversion of Pchlide by LPOR in the oligomers results in the oligomeric Chlide-LPOR-NADP^+^ complexes, which emit a fluorescence band peaking around 690 nm at 77 K. Then, NADP^+^ is replaced by NADPH and the oligomeric complexes are dissociated into dimeric forms. These processes are observed as a gradual shift (Shibata shift) of emission peaks from 690 nm to 680 nm at 77 K (Shibata, 1957; Smeller et al., 2003; Solymosi et al., 2007). In addition, some oligomeric Chlide-LPOR-NADP^+^ complexes exchange Chlide for Pchlide prior to the replacement of NADP^+^ to rapidly regenerate the oligomeric Pchlide-LPOR-NADPH complexes (Franck et al., 1999).

When dark-germinated seedlings are continuously illuminated, PLBs are transformed to the thylakoid membrane and etioplasts differentiate to chloroplasts (Kowalewska et al., 2016). Although PLBs greatly differ from the thylakoid membrane in morphology, the lipid compositions of these membranes are similar, with two galactolipids monogalactosyldiacylglycerol (MGDG) and digalactosyldiacylglycerol (DGDG) accounting for ∼80 mol% of total membrane lipids in both membranes (Fujii et al., 2019). In addition, sulfoquinovosyldiacylglycerol (SQDG) and phosphatidylglycerol (PG), which are anionic lipids with a negative charge in their polar head groups, make up the remaining 20 mol%. In the thylakoid membrane of chloroplasts, these lipids provide the fluid matrix for the photosynthetic complexes and prevent free diffusion of protons and other ions across the membrane. In addition, a number of lipid molecules are embedded in the structure of photosynthetic complexes including photosystem I and photosystem II and would play essential roles in the photochemical and electron transport reactions (Yoshihara and Kobayashi, 2022). Particularly, the necessity of PG for photosynthesis is evident as loss of the PG in the thylakoid membrane severely disrupt photosynthetic reactions in both cyanobacteria and plants (Kobayashi et al., 2016).

The importance of galactolipids in etioplasts was revealed by the analyses of *Arabidopsis thaliana* mutants deficient in MGDG or DGDG biosynthesis. A 36% decrease in MGDG content impaired Pchlide biosynthesis and the formation and oligomerization of the Pchlide-LPOR-NADPH ternary complex, while only slightly affecting the lattice structure of PLBs (Fujii et al., 2017). This is consistent with the data showing that MGDG mediates the oligomerization of the ternary complex in vitro (Gabruk et al., 2017). Similarly, an 80% decrease in DGDG content impaired the Pchlide biosynthesis and the formation of the ternary complex, although it did not affect the oligomerization of the ternary complex (Fujii et al., 2018). Moreover, the DGDG deficiency slowed the regeneration of the Pchlide–LPOR–NADPH complex from the Chlide–LPOR–NADP^+^ complex and severely disrupted the lattice structure of PLBs and the development of PTs. These data indicate a wide involvement of galactolipids in the membrane-associated processes during etioplast development. By contrast, roles of PG and SQDG in Pchlide accumulation, organization of the pigment-LPOR complexes, and internal membrane biogenesis during etioplast development remain unknown.

To address how PG and SQDG are involved in the processes of etioplast development in dark-germinated *A. thaliana*, we used the *pgp1-1* mutant, which is partially deficient in PG biosynthesis (Xu et al., 2002), and the *sqd1* and *sqd2-2* mutants, which both completely lack SQDG biosynthesis (Okazaki et al., 2009; Okazaki et al., 2013). The *pgp1-1* mutant carries a single amino acid substitution (P170S) in the major phosphatidylglycerophosphate synthase (PGP1) for the PG biosynthesis in plastids and mitochondria. The *pgp1-1* mutation decreases the PGP1 activity by 80% from the wild-type level, which results in a 30% reduction of total PG content and decreased Chl content and photosynthetic activity in leaves of light-grown plants (Xu et al., 2002). The *sqd1* (Okazaki et al., 2009) and *sqd2-2* (Okazaki et al., 2013) mutants have T-DNA insertions in the UDP-sulfoquinovose synthase gene (*SQD1*) and the SQDG synthase gene (*SQD2*), respectively, both of which are essential for SQDG biosynthesis. Although both *sqd1* and *sqd2-2* grew similar to wild type under nutrient-sufficient conditions, these mutants showed decreased photosynthetic activity when the PG content was decreased under phosphate-starved conditions. Furthermore, the *sqd1 pgp1-1* and *sqd2-2 pgp1-1* double mutants showed stronger impairments in photosynthesis and growth than the parental single mutants, indicating that PG and SQDG substitute each other (Yoshihara et al., 2021), as they share common features of anionic lipids (Yu and Benning, 2003). Of note, indistinguishable phenotypes between the *sqd1 pgp1-1* and *sqd2-2 pgp1-1* double mutants indicate that another anionic glycolipid glucuronosyldiacylglycerol (GlcADG), whose synthesis requires SQD2 but not SQD1, do not compensate for the loss of PG in *A. thaliana* (Okazaki et al., 2013; Yoshihara et al., 2021).

In this study, we investigated the membrane structures of etioplasts, the biosynthesis and accumulation of Pchlide, and the functions of the Pchlide-LPOR-NADPH complex in *pgp1-1*, *sqd1*, and the *sqd1 pgp1-1* double mutants. Some data were also obtained from *sqd2-2* and the *sqd2-2 pgp1-1* double mutant to corroborate the role of SQDG in etioplasts.

## RESULTS

### Changes in lipid composition in etiolated seedlings of the anionic lipid mutants

To reveal how the *pgp1-1* and *sqd1* mutations affect the lipid metabolism in etiolated seedlings, we determined the lipid composition in 4-d-old seedlings of wild type, *sqd1*, *pgp1-1*, and *sqd1 pgp1-1* grown under continuous darkness (Fig. 1). In *sqd1* and *sqd1 pgp1-1* mutants, no SQDG was detected, consistent with the previous report that the knockout mutation of *SQD1* causes complete loss of SQDG (Okazaki et al., 2009). Although the mean values of the proportion of PG was ∼25% lower in *pgp1-1* and *sqd1 pgp1-1* and 36% higher in *sqd1* than that in wild type, these differences were not statistically significant. When the total anionic lipid content (SQDG + PG) was compared between wild type and the mutants, only the *sqd1 pgp1-1* double mutant showed a significant decrease (Fig. S1A). Meanwhile, the proportion of both MGDG and DGDG was not significantly changed in all mutant lines (Fig. 1). In addition, there was no significant difference in the total amount of plastidic lipids (MGDG + DGDG + SQDG + PG) between the wild type and the mutants (Fig. S1B).

**Figure 1.**
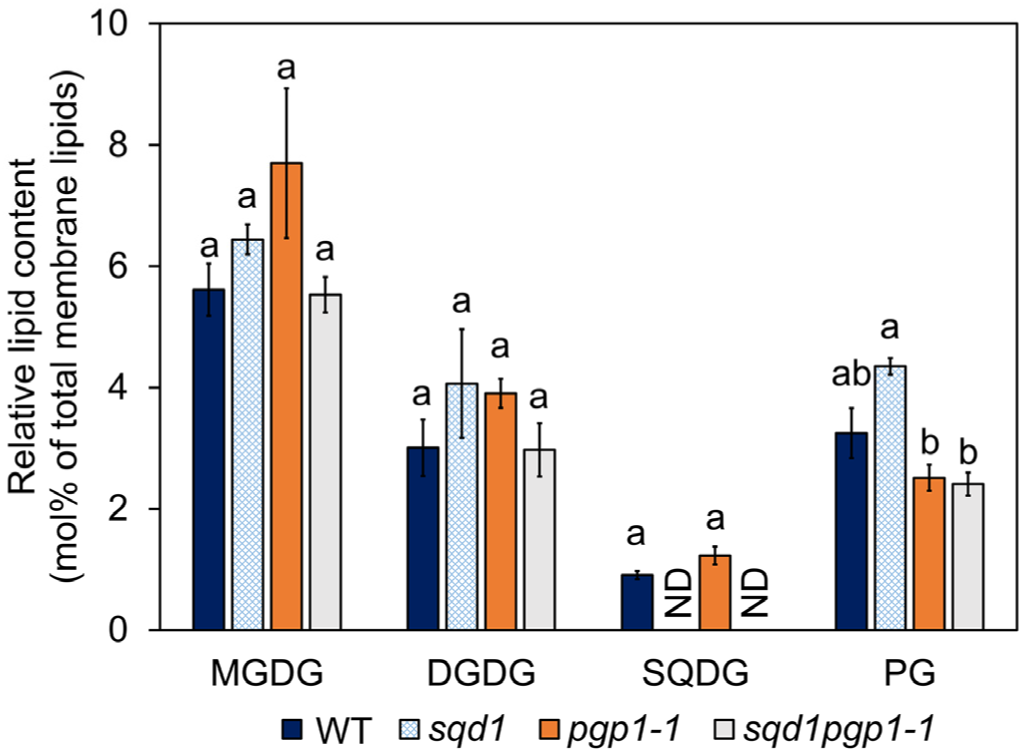
Proportion of major plastidic lipids in total membrane glycerolipids in 4-d-old etiolated seedlings of wild type and anionic lipid mutants. Data are means ± SE from three independent experiments. The lowercase letters indicate statistically significant differences between each line (p < 0.05, Tukey’s post hoc honestly significant difference test). MGDG, monogalactosyldiacylglycerol; DGDG, digalactosyldiacylglycerol; SQDG, sulfoquinovosyldiacylglycerol; PG, phosphatidylglycerol. ND, not detected.

In *pgp1-1* and *sqd1 pgp1-1*, fatty acid compositions of galactolipids but not anionic lipids were significantly changed as compared to those in wild type. In both mutants, proportions of 16:3 in MGDG and 16:0 in DGDG were increased with decreased proportions of 18:3 in both lipids (Fig. S2). Although some minor changes in fatty acid composition were observed in the mixture of other phospholipids (phosphatidylcholine + phosphatidylethanolamine + phosphatidylinositol) between each plant, the patterns of the changes were greatly different from those in galactolipids.

### Disordered membrane structures of etioplasts in anionic lipid mutants

To assess whether anionic lipids contribute to the membrane organization in etioplasts, we observed the ultrastructure of etioplasts in cotyledons of wild type and the anionic lipid mutants (Fig. 2). The wild type etioplasts developed highly regular lattice membrane structures of PLBs and long PTs connected to the PLBs (Fig. 2A-D). The membrane structures of the *sqd1* etioplasts were indistinguishable from those of wild type. By contrast, the PLB network was loosened in the *pgp1-1* etioplasts, which resulted in decreased membrane area in PLBs as compared with those in wild type and *sqd1* (Fig. 2E). Disorder of the PLB membrane structure was further enhanced in the etioplasts of the *sqd1 pgp1-1* double mutant. In the double mutant, some etioplasts showed loose aggregates of filamentous membrane tubules in stromal region, although some showed electron-dense aggregates of small membranes. In addition, the *sqd1 pgp1-1* double mutant frequently showed etioplast sections that had almost no PLB and PT membranes (Fig. 2F), which were rarely observed in wild-type and single mutants. Meanwhile, the double mutant showed no significant differences in the circularity (Fig. 2G) and size of etioplasts (Fig. 2H) compared with the wild type and the single mutants. We also observed ultrastructures of etioplasts in *sqd2-2* and *sqd2-2 pgp1-1* mutants (Fig. S3). The *sqd2-2 pgp1-1* etioplasts showed disordered PLB structures similar to *sqd1 pgp1-1*, whereas the *sqd2-2* etioplasts were comparable to those in wild type and *sqd1*.

**Figure 2.**
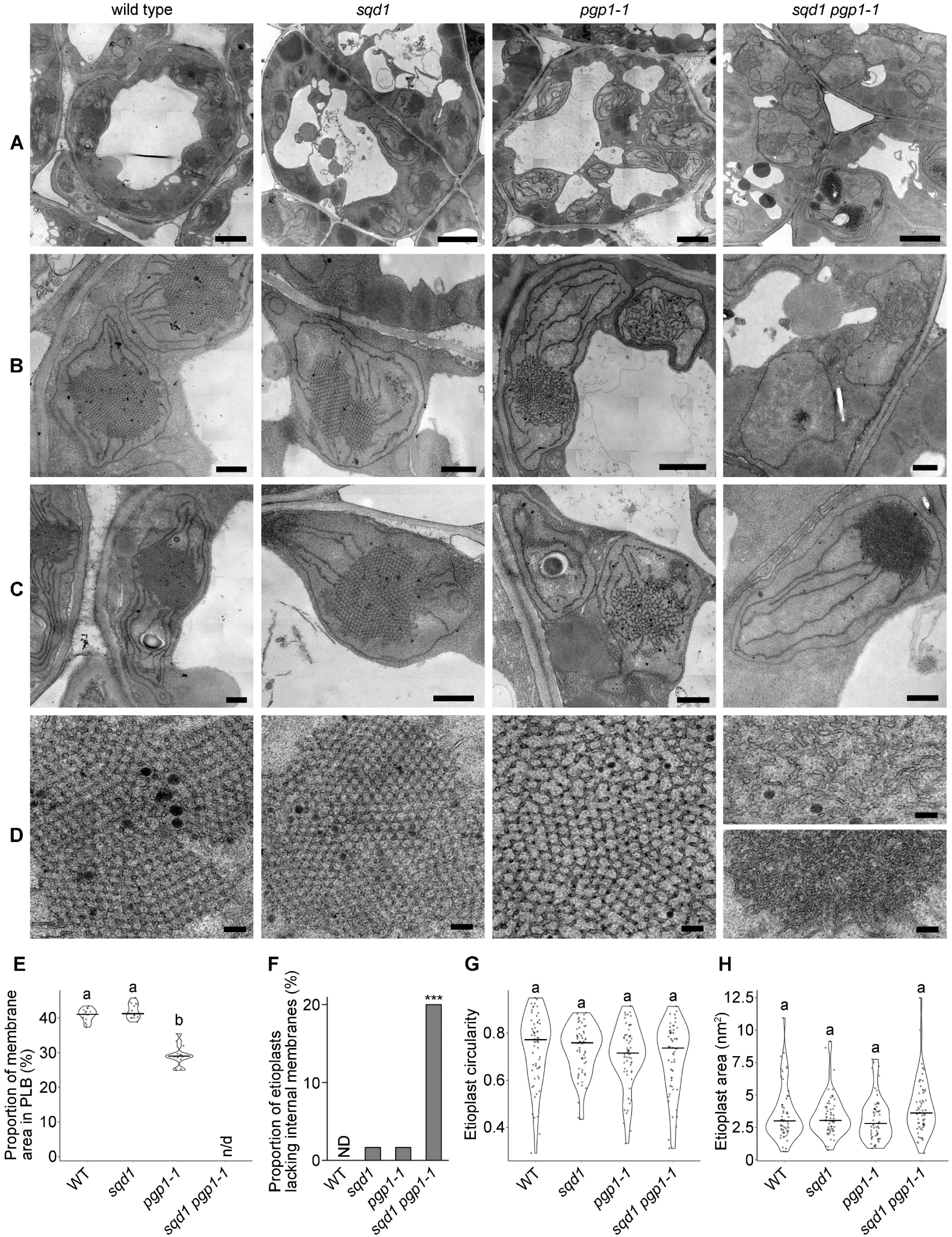
Structure analysis of cotyledon etioplasts in 4-d-old etiolated seedlings of wild type and anionic lipid mutants. Ultrastructures of cells (A), etioplasts (B and C), and internal membranes of etioplasts (D) in wild type, *sqd1*, *pgp1-1*, and *sqd1 pgp1-1*. In D, two different types of internal membrane structures were shown for the *sqd1 pgp1-1* etioplasts. Bars = 2 μm in A, 500 nm in B and C, and 100 nm in D. E, Proportion of membrane area in PLB. F, Proportion of etioplasts lacking internal membranes. G, Etioplast circularity. H, Area of etioplasts. In E, G, and H, grey dots show the raw data, the width of the violin plots represents the density of the data, and horizontal bars indicate median values. The lowercase letters indicate statistically significant differences between each line (p < 0.05, Tukey’s post hoc honestly significant difference test). Asterisks (***) in F indicate a significant difference from the wild type (P < 0.001, Pearson’s chi-square test). n/d, not determined. ND, not detected.

Because the LPOR protein is involved in the development of PLBs (Sperling et al., 1998; Franck et al., 2000), we determined LPOR levels in etiolated wild type and the anionic lipid mutants by using polyclonal anti-LPOR antibodies that react with all Arabidopsis LPOR isoforms (Masuda et al., 2003). The *sqd1 pgp1-1* and *sqd2-2 pgp1-1* double mutants showed ∼60% decreases in the LPOR level compared with the wild type (Fig. 3). By contrast, no significant change in LPOR levels was detected in the *sqd1*, *sqd2-2*, and *pgp1-1* mutants.

**Figure 3.**
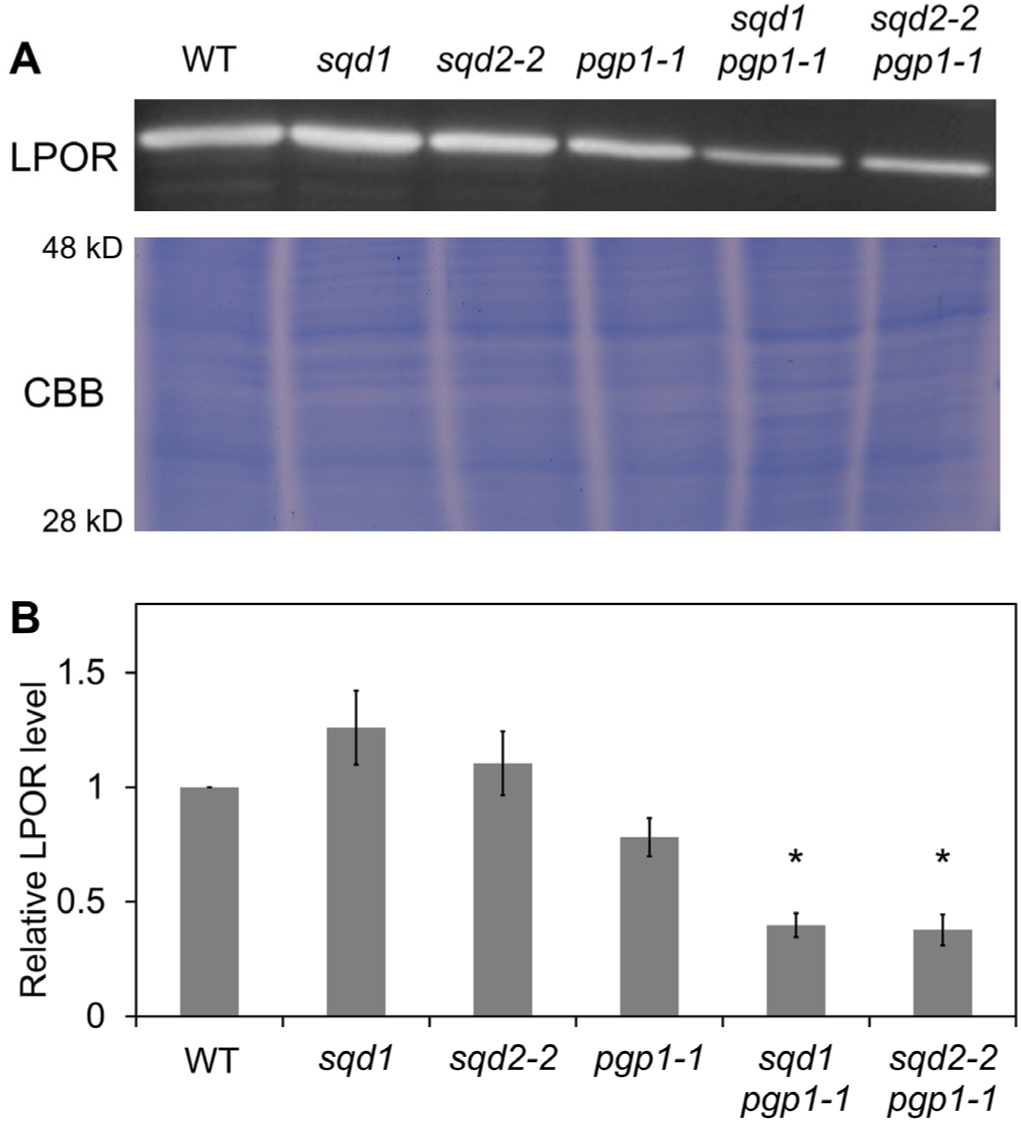
LPOR levels in 4-d-old etiolated seedlings of wild type and anionic lipid mutants. A, Immunodetection of total LPOR proteins (∼37 kDa) in 4-d-old etiolated seedlings of wild type and anionic lipid mutants. Coomassie brilliant blue staining of the same protein samples are shown as a loading control. B, LPOR protein levels in the anionic lipid mutants relative to the wild-type level. Data are means ± SE from three independent experiments. Asterisks indicate statistically significant differences from the wild-type level (p < 0.05, Student’s t-test).

### Decreased Pchlide content in etiolated seedlings of anionic lipid mutants

To determine the role of PG and SQDG in Pchlide accumulation, we measured Pchlide content in 4-d-old etiolated wild type and the anionic lipid mutants (Fig. 4). Total Pchlide content in *sqd1* and *sqd2-2* was similar to and slightly higher than that in the wild type, respectively. By contrast, total Pchlide content decreased by 41% in *pgp1-1* and by ∼70% in the *sqd1 pgp1-1* and *sqd2-2 pgp1-1* mutants compared to the wild-type level. The amounts of Pchlide remaining after a 0.7 ms flash, which corresponds to nonphotoactive Pchlide, also decreased by 52% in *pgp1-1* and by ∼70% in both double mutants from the wild-type level. As a result, the ratio of nonphotoactive Pchlide to total Pchlide was not notably changed between wild type and these mutants (Fig. 4B).

**Figure 4.**
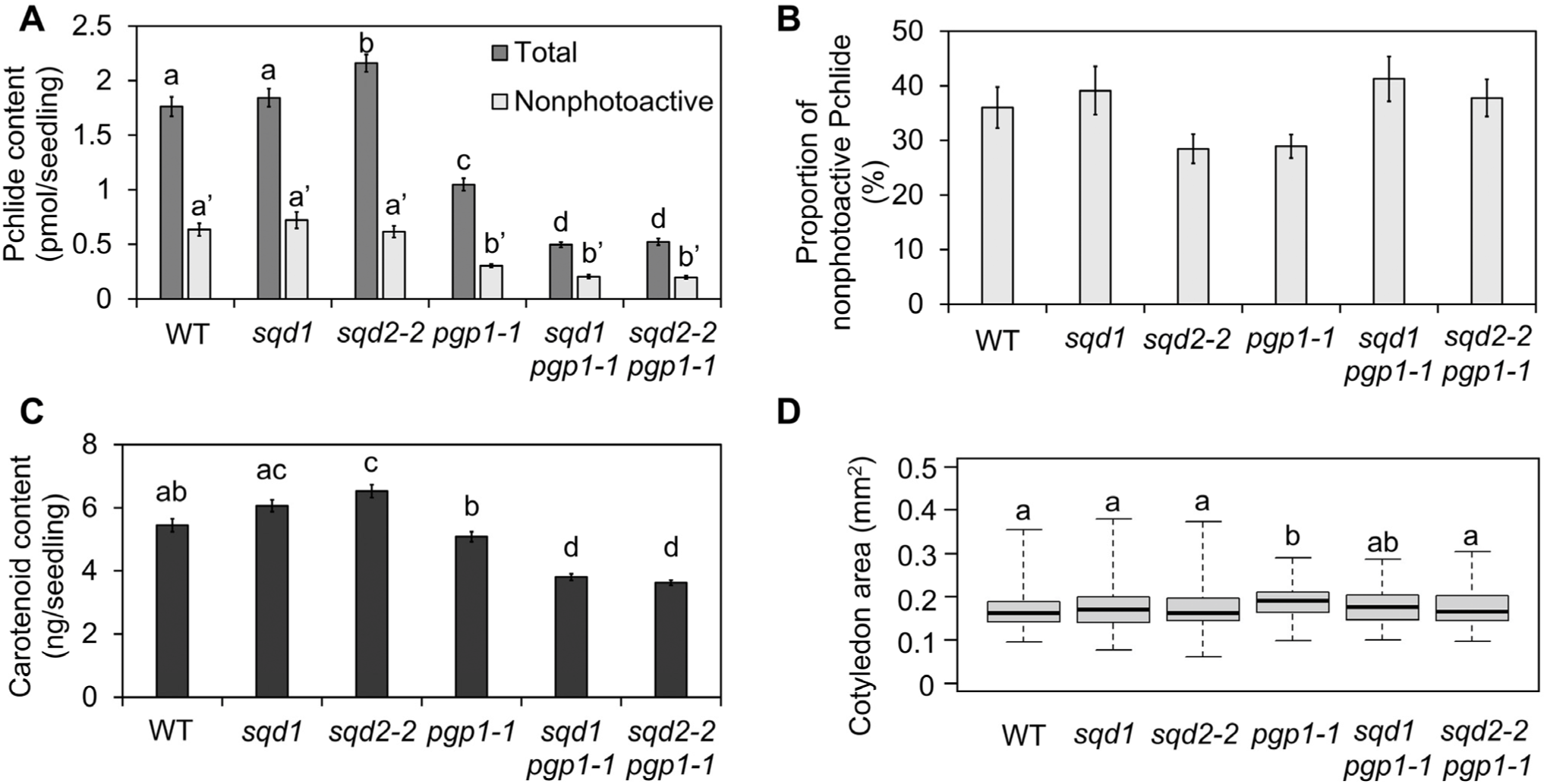
Pigment content in 4-d-old etiolated seedlings of wild type and anionic lipid mutants. A, Content of total and nonphotoactive Pchlides, which were extracted before and after flash treatment, respectively. Data are means ± SE from 16 independent experiments. The lowercase letters (a-d for total Pchlide and a’ and b’ for nonphotoactive Pchlide) indicate statistically significant differences between each line (p < 0.05, Tukey’s post hoc honestly significant difference test). B, Proportion of nonphotoactive Pchlide to total Pchlide. Data are represented as means ± SE from 16 independent experiments. No significant difference was observed among all plants (P > 0.05, Welch’s t test). C, Total carotenoid content. Data are means ± SE from 5 independent experiments. The lowercase letters indicate statistically significant differences between each line (p < 0.05, Tukey’s post hoc honestly significant difference test). D, Size of 4-d-old etiolated cotyledons. Data are represented as means ± SE from 196 cotyledons. The horizontal line in each box represents the median value of the distribution. The top and bottom of each box represent the upper and lower quartiles, respectively. The whiskers represent the range. The lowercase letters indicate statistically significant differences between each line (p < 0.05, Tukey’s post hoc honestly significant difference test).

We also determined total carotenoid levels in etiolated seedlings. The *sqd1 pgp1-1* and *sqd2-2 pgp1-1* mutants but not *sqd1*, *sqd2-2*, and *pgp1-1* single mutants showed slight reductions in carotenoid content compared with the wild type (Fig. 4C). By contrast, the cotyledon size of the double mutants was comparable to that of wild type (Fig. 4D), suggesting that the seedling growth was not impaired in the mutants.

### Impairment of the Pchlide biosynthesis pathway in anionic lipid mutants

To examine which step of the Pchlide biosynthesis pathway is impaired in anionic lipid mutants, we treated etiolated seedlings with ALA in darkness for 24 h to bypass the rate-limiting step of the Pchlide biosynthesis pathway and determined the accumulation levels of Pchlide and the porphyrin intermediates (Proto IX, Mg-Proto IX, and Mg-Proto IX ME). In wild type and *sqd1*, the ALA treatment did not cause remarkable accumulation of all three porphyrin intermediates (Fig. 5A). By contrast, *pgp1-1* and *sqd1 pgp1-1* showed strong accumulation of Proto IX without noticeable increases in Mg-Proto IX and Mg-Proto IX ME content, which suggests that the conversion of Proto IX to Mg-Proto IX is particularly impaired in these mutants. Furthermore, the *sqd1 pgp1-1* mutant showed lower Pchlide accumulation even in the presence of ALA. The *sqd2-2 pgp1-1* mutant also showed specific accumulation of Proto IX along with impaired accumulation of Pchlide in the presence of ALA (Fig. S4).

**Figure 5.**
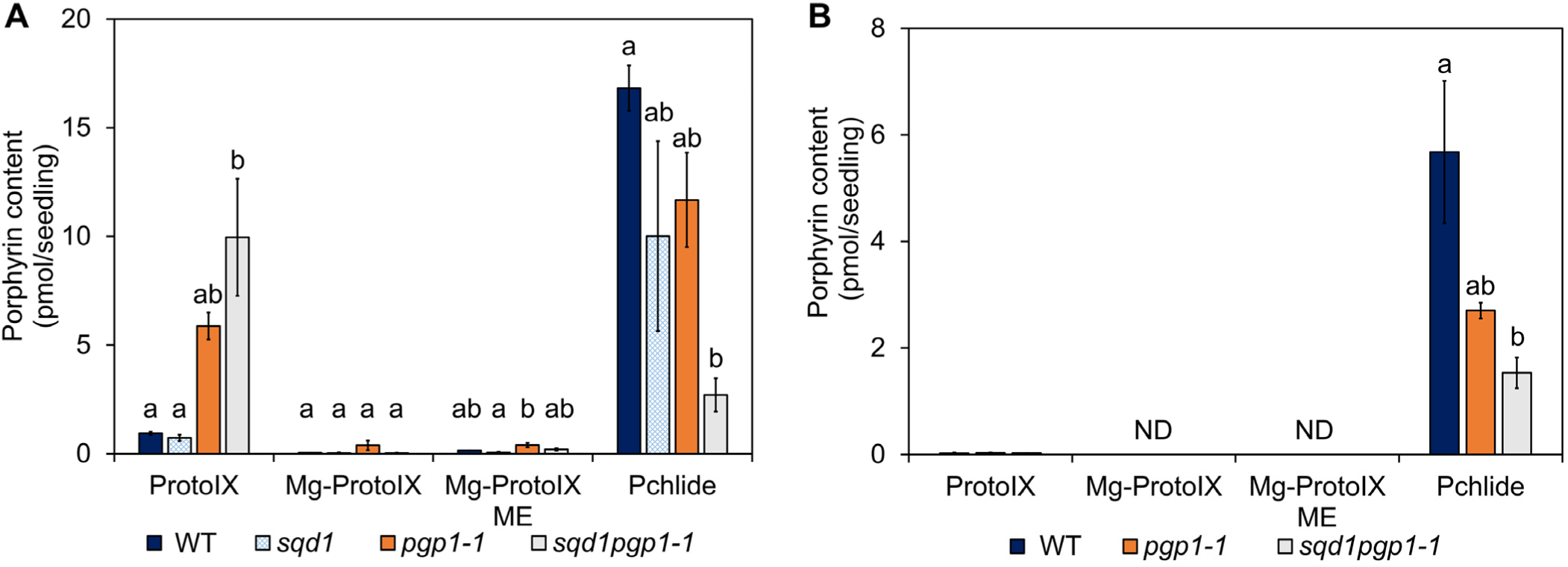
Altered Pchlide metabolism in anionic lipid mutants compared with wild type. Accumulation levels of porphyrin pigments in 4-d-old etiolated seedlings treated with (A) or without (B) 10 mM ALA for 24 h in the dark. Data are represented as means ± SE from four (A) and three (B) independent experiments. The lowercase letters indicate statistically significant differences between each line (p < 0.05, Tukey’s post hoc honestly significant difference test). ND, Not detected.

Then we tested whether the *pgp1-1* and *sqd1 pgp1-1* mutants accumulate the porphyrin intermediates even in the absence of exogenous ALA. As in the wild type, only negligible amount of Proto IX was accumulated in both *pgp1-1* and *sqd1 pgp1-1*, with Mg-Proto IX and Mg-Proto IX ME being undetectable (Fig. 5B).

To ascertain whether the impaired conversion of Proto IX to Mg-Proto IX in *pgp1-1* and *sqd1 pgp1-1* is caused at the transcription level, we investigated mRNA levels of *CHLD, CHLH,* and *CHLI1* genes in etiolated seedlings. *CHLD* and *CHLH* encode the D and the H subunit of Mg-Chelatase, respectively, whereas *CHLI1* encodes the major isoform of the I subunits. We also examined the mRNA level of *GUN4*, which encodes the GUN4 protein required for Mg-chelatase activity (Tanaka et al., 2011). Excepting *GUN4* in *sqd1 pgp1-1*, no significant difference in mRNA levels was observed between wild type and the anionic lipid mutants (Fig. S5).

### Plastidic anionic lipids affect the behavior of Chlide-LPOR complexes after photoconversion

To examine how the deficiency of anionic lipids affects the activity and the behavior of the LPOR-pigment complexes, we examined in situ low temperature fluorescence spectra from etiolated cotyledons of wild type and anionic lipid mutants. At 77K, two emission bands peaking at 634 and 657 nm, which were attributed to nonphotoactive and photoactive Pchlide, respectively, were observed in wild type and all anionic lipid mutants (Fig. 6A). With a 0.7-ms flash treatment, the emission band around 657 nm disappeared in all plants and a new band emerged around 695 nm, which originates from Chlide in oligomeric Chlide-LPOR-NADP^+^ complexes (Fig. 6B). The data demonstrate that the instantaneous photoconversion of Pchlide to Chlide by LPOR efficiently occurred in the anionic lipid mutants as well as wild type.

**Figure 6.**
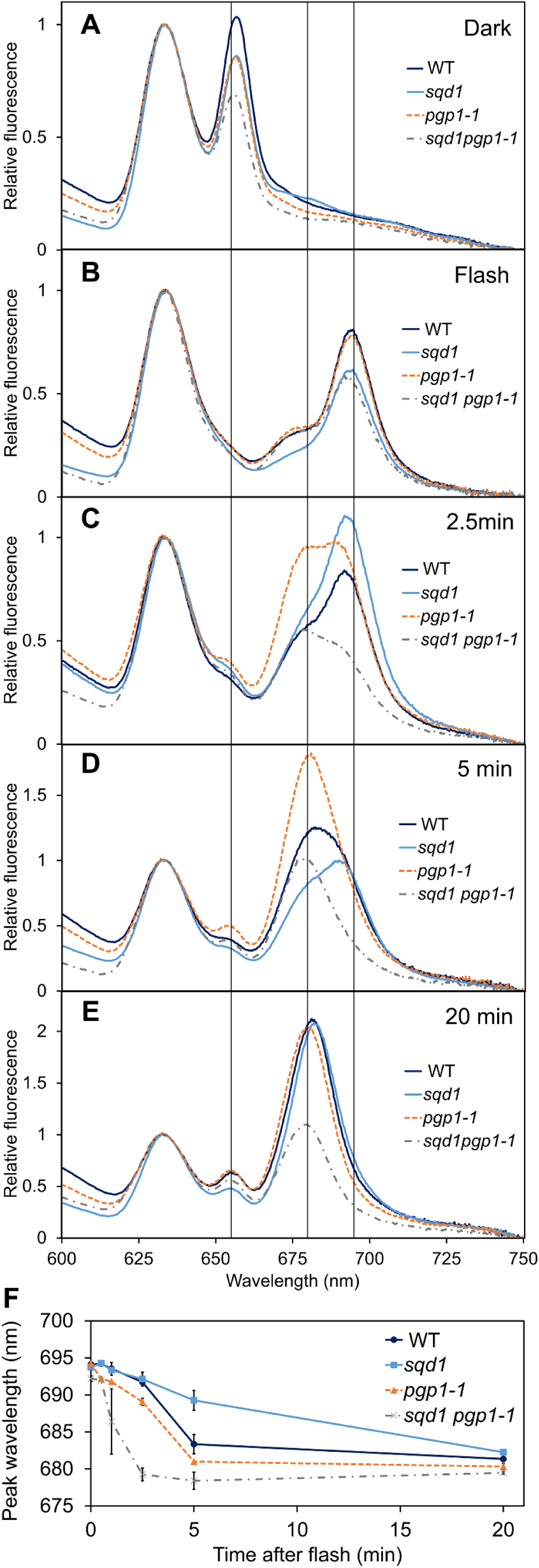
In situ 77K fluorescence spectra in 4-d-old etiolated seedlings of the wild type and anionic lipid mutants. A and B, Fluorescence spectra in cotyledons frozen in liquid nitrogen before (A) and immediately after (B) the flash treatment. C to F, Fluorescence spectra in cotyledons frozen after dark incubation for 2.5 min (C), 5 min (D), and 20 min (E) after the flash treatment. Representative data from three independent experiments are shown. Vertical lines are on 655 nm, 680 nm, and 695 nm. F, Overview of the Shibata shift after the flash treatment. Data are represented as means ± SE from three independent experiments.

After photoconversion, a gradual shift (Shibata shift) of the Chlide fluorescence was observed in etiolated cotyledons during dark incubation, which reflects the dissociation process of the oligomeric Chlide-LPOR-NADP^+^ complex (Shibata, 1957; Smeller et al., 2003; Solymosi et al., 2007). At 2.5 min after the flash, wild type and *sqd1* showed a shoulder band near 680 nm in addition to the peak near 695 nm (Fig. 6C). By contrast, *pgp1-1* and *sqd1 pgp1-1* showed a strong fluorescence around 680 nm already at 2.5 min after the flash, suggesting a faster shift of the Chlide fluorescence in these mutants. At 5 min after the flash, *sqd1* still showed the Chlide fluorescence peaking near 690 nm, although the wild type showed a peak at 683 nm (Fig. 6D). In *pgp1-1* and *sqd1 pgp1-1*, Chlide fluorescence peaks were observed at 681 nm and 678 nm, respectively, at 5 min. At 20 min after the flash, all plants showed Chlide fluorescence peaks around 680 nm (Fig. 6E). The quantification of the peak wavelengths of Chlide fluorescence demonstrates that the shift proceeded most rapidly in *sqd1 pgp1-1*, followed by *pgp1-1*, wild type, and *sqd1* (Fig. 6F). The *sqd2-2* and the *sqd2-2 pgp1-1* mutant also showed slower and faster shifts similar to *sqd1* and *sqd1 pgp1-1*, respectively (Fig. S6), confirming the effects of SQDG deficiency on the dissociation process of the Chlide-LPOR oligomers.

## DISCUSSION

### PG biosynthesis affects various processes of etioplast development

The *pgp1-1* mutant carries a point mutation in the *PGP1* gene and has an 80% reduction in PGP1 activity, which results in a 30% reduction in PG content in light-grown leaves (Xu et al., 2002). The small difference in the PG content between *pgp1-1* and wild type can be partly explained by the activity of PGP2, which is the minor isoform of phosphatidylglycerophosphate synthase localized to the endoplasmic reticulum (ER) (Tanoue et al., 2014), in addition to the remaining PGP1 activity in this mutant. Although the mean value of PG content was 23% lower in etiolated *pgp1-1* seedlings than the wild type, the difference was not statistically significant (Fig. 1). The data indicate that the effect of the *pgp1-1* mutation on total PG content was small or negligible in etiolated seedlings. By contrast, the *pgp1-1* mutation had strong impacts on the multiple processes in etioplasts including formation of the regular PLB structure (Fig. 2), Pchlide biosynthesis (Fig. 4 and 5), and LPOR-pigment dynamics (Fig. 6). Therefore, not the change in total PG content in the cell but the local disruption of the PG biosynthesis in etioplasts would strongly affect these processes in *pgp1-1*. Although PGP1 is targeted to mitochondria in addition to plastids (Babiychuk et al., 2003), it is unlikely that the disruption of mitochondrial PG biosynthesis in *pgp1-1* strongly affects the processes in etioplasts, because even the knockout mutants of PGP1, which have severe disruptions in chloroplast development with a seedling lethal phenotype, have functional mitochondria in leaves (Hagio et al., 2002; Babiychuk et al., 2003).

Previous studies revealed that deficiencies of galactolipids also disrupt the PLB structure, Pchlide biosynthesis, and LPOR-pigment dynamics (Fujii et al., 2017; Fujii et al., 2018). In *pgp1-1*, the contents of galactolipids (MGDG + DGDG), anionic lipids (SQDG + PG), and total plastidic lipids (MGDG + DGDG + SQDG + PG) were comparable to those in wild type (Fig. 1 and Fig. S1), so the *pgp1-1* mutation affected etioplast development without changing the global lipid levels in etioplasts. In contrast, the *pgp1-1* mutation significantly changed the fatty acid compositions of MGDG and DGDG, although it did not strongly affect those of SQDG, PG, and other phospholipids (Fig. S2). It is unknown how the *pgp1-1* mutation specifically affects the fatty acid compositions of MGDG and DGDG. In plant cells, galactolipids are synthesized via two distinct pathways; the plastid pathway completed within plastids and the ER pathway via phospholipid biosynthesis in the ER (Li-Beisson et al., 2013). Because 16:3-containing MGDG, which is exclusively synthesized via the plastid pathway, was increased by the *pgp1-1* mutation, the plastid pathway may be enhanced by the decreased PGP1 activity or consequent impairments in etioplast development. We cannot exclude that the changes in fatty acid compositions of galactolipids by *pgp1-1* caused etioplast disruptions. Nevertheless, the *sqd1 pgp1-1* double mutant showed stronger disruptions in etioplast development than *pgp1-1* despite the similar fatty acid compositions of galactolipids between these mutants. Thus, we assume that the lack of anionic lipids would be the main cause of the perturbed etioplast development in *pgp1-1* and the double mutants.

In contrast to the strong effects of the partial limitation of PG biosynthesis on the etioplast development, complete lack of SQDG did not notably change the PLB structure (Fig. 2 and Fig. S3) and Pchlide biosynthesis (Fig. 4 and 5). The data indicate that SQDG is not essential for these processes in Arabidopsis. In *sqd1*, total anionic lipid content was maintained to the wild-type level (Fig. S1A), so PG may function to compensate the loss of SQDG in the *sqd1* etioplasts. Of note, the additional mutation of *sqd1* in *pgp1-1*, which resulted in a significant decrease in total anionic lipid content, strongly enhanced the defects in PLB formation and Pchlide biosynthesis, suggesting the importance of SQDG when the PG biosynthesis is limited. We confirmed that the *sqd2-2* mutation affected the etioplast functions similar to *sqd1*, suggesting that GlcADG, which is abolished in *sqd2-2* but not in *sqd1*, has no important function in etioplasts at least under nutrient-sufficient conditions. As suggested in the process of chloroplast development (Yu and Benning, 2003; Yoshihara et al., 2021), SQDG in etioplasts would play a role in maintaining total anionic lipid constant, whereas PG has specific functions that cannot be complemented by SQDG even during etioplast development.

### Anionic lipids are essential for the internal membrane formation in etioplasts

The PLB lattice structure was disrupted by the *pgp1-1* mutation. Nguyen et al. (2021) reported that Arabidopsis LPORs require PG and MGDG in addition to Pchlide and NADPH to form in vitro helical tubes that fluorometrically resemble isolated PLBs. Because LPOR is a direct membrane-binding and membrane-remodeling enzyme, an interaction between LPOR and PG may be important for PLBs to form the regular lattice.

The *sqd1 pgp1-1* double mutant showed impaired formation of PTs in addition to the severely disrupted PLB structures (Fig. 2). It was reported that the *dgd1* mutation, which caused an 80% decrease in DGDG content, also impaired the development of PTs in addition to disordering the PLB lattice (Fujii et al., 2018). Because DGDG is the second most abundant lipid in etioplasts, the substantial decrease in this lipid greatly alters the ratio of the non-bilayer-forming MGDG to bilayer-forming lipids (DGDG + SQDG + PG) (Jouhet, 2013) and may also cause a deficiency of lipid molecules to extend the lamellar membranes in etioplasts. By contrast, the *sqd1 pgp1-1* double mutations did not largely affect the abundance of total plastidic lipids (Fig. S1B) and the ratio of the non-bilayer-to-bilayer-forming lipids in etioplasts (Fig. 1). Thus, not a lack of total bilayer lipids but specific loss of the anionic lipids would perturb PT formation in *sqd1 pgp1-1* etioplasts. Another possibility is that the decreased abundance of LPORs in the double mutant results in the decreased PT formation in etioplasts (Fig. 3). However, considering that antisense RNA inhibition of *LPOR* expression in *A. thaliana* did not impair the formation PTs (Franck et al., 2000), the effect of decreased abundance of LPORs in *sqd1 pgp1-1* on the PT formation would be marginal.

### Anionic lipids are essential for the Pchlide biosynthesis pathway

In the *pgp1-1* mutant, Pchlide content significantly decreased, which was further enhanced in the *sqd1 pgp1-1* and *sqd2-2 pgp1-1* double mutants (Fig. 4). ALA feeding experiments showed specific accumulation of Proto IX in *pgp1-1* and the double mutants (Fig. 5A and Fig. S4), indicating that the Mg insertion into Proto IX by Mg chelatase was specifically impaired by lack of anionic lipids. This result contrasts with the reports that the deficiencies of MGDG and DGDG both caused high accumulation of Mg-Proto IX with ALA feeding to etiolated seedlings (Fujii et al., 2017; Fujii et al., 2018). Thus, anionic lipids play a crucial role in the Mg insertion step by Mg chelatase, whereas galactolipids are required for the Mg-Proto IX metabolism by MgMT. Of note, deficiencies of galactolipids also caused the accumulation of Proto IX and Mg-Proto IX ME with ALA feeding, suggesting a broad effect of galactolipids on the membrane-associated steps of the Pchlide biosynthesis pathway (Fujii et al., 2019). By contrast, anionic lipids may be more specifically involved in the Mg insertion step.

Because the mRNA expression of the genes involved in the Mg insertion step was not suppressed in *pgp1-1* (Fig. S5), transcriptional regulation would be irrelevant to this impairment. Nevertheless, we cannot exclude the possibility that the enzymes involved in this step are not sufficiently accumulated or recruited to the etioplast membranes, in addition to the possibility that the Mg-chelatase requires anionic lipids to exert its activity. Decreased Chl biosynthesis by loss of PG biosynthesis was also observed in the cyanobacterium *Synechocystis* sp. PCC 6803 (Kopečná et al., 2015). Although the requirement of PG for the Mg insertion step remains unknown in cyanobacteria, these data demonstrate the importance of PG for the Chl biosynthesis pathway across cyanobacteria and plants.

High accumulation of Proto IX in *pgp1-1* and *sqd1 pgp1-1* by ALA treatment suggest that the biosynthetic pathway from ALA to Proto IX is functional in this mutant (Fig. 5A). Meanwhile, in the absence of ALA, neither *pgp1-1* nor *sqd1 pgp1-1* showed excess accumulation of Proto IX and other porphyrin intermediates despite the impaired Pchlide biosynthesis (Fig. 5B). These data imply that, in *pgp1-1* and *sqd1 pgp1-1*, Pchlide biosynthesis is suppressed at the pathway preceding the ALA formation, as similarly observed in MGDG and DGDG-deficient *A. thaliana* mutants (Fujii et al., 2017; Fujii et al., 2018).

### Disruptions of PG biosynthesis and SQDG biosynthesis oppositely affect the dynamics of Chlide-LPOR complex after photoconversion

In *pgp1-1* and *sqd1 pgp1-1*, the ratio of photoactive Pchlide to total Pchlide was similar to that in the wild type (Fig. 4B). Moreover, in these mutants, all photoactive Pchlide was instantaneously converted to Chlide with a flash (Fig. 6B). Although PG was shown to increase the affinity of LPORs towards NADPH in vitro (Gabruk et al., 2017), our data indicate that PG and SQDG are not required for the formation of photoactive Pchlide-LPOR complexes in vivo or that the decreased amounts of PG in *pgp1-1* and *sqd1 pgp1-1* are still sufficient for the photoactive complex formation. In addition, unlike the MGDG-deficient *A. thaliana* plants (Fujii et al., 2017), *pgp1-1* and *sqd1 pgp1-1* showed no remarkable fluorescence emission from the dimeric forms of the photoactive complex around 645 nm at 77 K (Fig. 6A), suggesting that the partial deficiency of PG and complete loss of SQDG do not affect the oligomerization of the photoactive complexes.

After photoconversion of Pchlide to Chlide, low temperature fluorescence from oligomeric Chlide-LPOR complexes around 690 nm shifts to shorter wavelength (∼680 nm) during dark incubation, which reflects dissociation of the oligomeric complexes and the replacement of NADP^+^ with NADPH in the complex (Smeller et al., 2003; Solymosi et al., 2007). As compared with wild type, *pgp1-1* and *sqd1* showed faster and slower transitions of Chlide fluorescence, respectively (Fig. 6F), suggesting that the partial disruption of PG biosynthesis accelerates the dissociation of the oligomeric Chlide-LPOR complexes, whereas the complete loss of SQDG delays it. Surprisingly, the *sqd1 pgp1-1* double mutant showed even a faster shift of Chlide fluorescence than the *pgp1-1* single mutant. A compensatory increase in PG content in response to the loss of SQDG has been widely observed in plants (Yu and Benning, 2003), algae (Sato et al., 1995), and cyanobacteria (Güler et al., 1996; Aoki et al., 2004; Endo et al., 2016). In fact, the mean value of the relative PG content was 1.4-fold higher in *sqd1* than wild type, although the difference was not statistically significant (Fig. 1). We hypothesize that PG has an effect to prevent the dissociation of the oligomeric Chlide-LPOR complex, and thus a compensatory increase of the PG content in *sqd1* may stabilize the oligomeric Chlide-LPOR complexes, whereas the partial PG deficiency in *pgp1-1* may destabilize them. This assumption is consistent with the rapid dissociation of the Chlide-LPOR oligomers in *sqd1 pgp1-1*, whose PG content was not increased in response to the loss of SQDG. Because the PLB lattice structure was substantially loosened in *pgp1-1* and *sqd1 pgp1-1* (Fig. 2), the disorder of the membrane structure might affect the behavior of the Chlide-LPOR complexes. However, the *dgd1-1* mutant also showed disordered PLB lattice, although this mutant had retarded dissociation of the Chlide-LPOR oligomers (Fujii et al., 2018). Therefore, not the global change in the PLB structure but the loss of the specific interaction between anionic lipids and Chlide-LPOR complexes may affect the behavior of the Chlide-LPOR oligomers during the Shibata shift.

## MATERIALS AND METHODS

### Plant materials and growth conditions

The wild type, *sqd1* (Okazaki et al., 2009), *sqd2-2* (Okazaki et al., 2013), *pgp1-1* (Xu et al., 2002), *sqd1 pgp1-1* and *sqd2-2 pgp1-1* (Yoshihara et al., 2021) were the Columbia ecotype of *Arabidopsis thaliana*.

Surface-sterilized seeds on half strength Murashige and Skoog (MS) medium (adjusted to pH 5.7 with KOH) containing 1% (w/v) sucrose solidified with 0.8% (w/v) agar were cold-treated at 4℃ for 3 or 4 d in the dark. All plants were grown at 23℃ in a growth chamber. The cold-treated seeds were illuminated with white light (∼ 90 μmol photons m^-2^ s^-1^) for 4 h to synchronize germination and then germinated in the dark. Etiolated seedlings were sampled under dim green light unless stated otherwise.

### Lipid analysis

Total lipids were extracted from seedlings crushed in liquid nitrogen according to Bligh and Dyer (1959) and were separated by one dimensional thin layer chromatography as described (Sato et al., 2020) with a modified solvent system of acetone:toluene:methanol:water (40:15:5:2, v/v/v/v). After visualization with 0.01% (w/v) primuline in 80% (v/v) acetone under UV light, MGDG, DGDG, SQDG, PG, and a mixture of other phospholipids (phosphatidylcholine, phosphatidylethanolamine, and phosphatidylinositol) were scraped from silica gel plates. Fatty acids in each lipid fraction were methyl esterified by incubation in 1 M HCl in methanol at 85°C for 2 h and quantified by gas chromatography (GC-2014; Shimadzu; http://www.shimadzu.com/) with myristic acid as an internal standard.

### Transmission electron microscopy analysis

Fixation of etiolated cotyledons, sectioning of the fixed samples, staining of the ultrathin sections with uranyl acetate and lead citrate, and observation of the samples by transmission electron microscopy were performed as described previously (Fujii et al., 2017). Quantitative analysis of etioplast ultrastructure was performed with the ImageJ variant software Fiji (Schindelin et al., 2012). The proportion of membrane area in PLBs was determined by measuring the total membrane area in a 0.09 μm^2^ area of a PLB. To calculate the frequency of internal membrane-lacking etioplasts in ultrathin sections, we defined etioplasts with a membrane area less than 2% of the total area as “internal membrane-lacking etioplasts” and counted the number of them out of 60 etioplasts for each line. The circularity index of etioplasts was calculated as follows: 4 × ᴨ × (area of etioplast) / (perimeter of etioplast)^2^. Violin plots of the results were produced using the ggplot2 package in R program (Wickham, 2009).

### Immunoblot analysis

Extraction, quantification, and separation of total proteins from etiolated seedlings as well as detection and quantification of total LPOR proteins were performed as described previously (Fujii et al., 2017), with some modifications. Total protein content was determined using a protein assay reagent (XL-Bradford, Pharma Foods International, Kyoto) and 10 μg of total proteins were subjected to SDS-PAGE. The anti-LPOR antibody, which react with all Arabidopsis LPOR isoforms (Masuda et al., 2003), was used as a primary antibody. Goat anti-rabbit IgG antibody conjugated with horseradish peroxidase (SA00001-2, Proteintech) was secondarily reacted with the anti-LPOR antibody and detected using a chemiluminescence reagent (Immobilon; Merck Millipore) and an imager (LuminoGraph I; ATTO). For the loading control, total proteins separated by SDS-PAGE were stained with Coomassie brilliant blue.

### Determination of pigment content

To determine steady-state levels of Pchilde and carotenoids in 4-d-old etiolated seedlings, pigments were extracted in 80% (v/v) acetone and analyzed as described (Fujii et al., 2017). In brief, Pchlide content was determined by measuring the fluorescence of the extract at 634 nm under 433-nm excitation with a spectrofluorometer (RF-5300PC; Shimadzu) and a Pchlide standard of known concentration. The amount of nonphotoactive Pchlide was determined by illuminating intact seedlings with a 0.7-ms single flash from an electronic flash equipment (PZ42X, Sunpak) before pigment extraction. Carotenoid content was determined with the V-730 BIO spectrophotometer (JASCO) according to the following formula: 5.05 × absorbance at 470 nm (μg carotenoids mL^−1^; Lichtenthaler, 1987).

To analyze Pchlide synthesizing activity, 4-d-old etiolated seedlings were incubated in the dark in a solution containing 10 mM MES-KOH (pH 5.7) and 5 mM MgCl_2_, with or without 10 mM ALA, at 23°C for 24 h. Then porphyrin pigments were extracted from the seedlings in N, N-dimethylformamide and quantified by HPLC as described (Fujii et al., 2017). The HPLC system consists of L-6200 and L-6000 pumps (Hitachi), an injector with a 20-μL sample loop (LC-organizer, Hitachi), a L-column2 C8 guard column (5 µm, 4.6 × 10 mm; Chemicals Evaluation and Research Institute), a reverse-phase C8 column (Symmetry C8 column, 100 Å, 3.5 µm, 4.6 × 150 mm; Waters) and detected by using an FP-4025 spectrofluorometric detector (JASCO).

### Quantitative reverse transcription-PCR analysis

Total RNA was extracted by using the NucleoSpin RNA Plant (MACHEREY-NAGEL), followed by genomic DNA digestion and reverse transcription with use of the ReverTra Ace qPCR RT Master Mix with gDNA Remover kit (TOYOBO). Complementary DNA was amplified by thermal cycling consisting of an initial denaturation step at 95°C for 60 s followed by 40 cycle of 15 s at 95°C and 45 s at 60°C with use of the Thunderbird SYBR qPCR Mix (TOYOBO) and 200 nM gene-specific primers (Supplemental Table S1). The thermal cycling and signal detection were performed in duplicate by use of StepOne Real-Time PCR System (Applied Biosystems). The relative abundance of all transcripts amplified was normalized to the means of constitutive expression level of *ACTIN8* and *UBIQUITIN11* according to Pfaffl (2001).

### In situ low temperature fluorescence spectroscopy

Fluorescence emission spectra from Pchlide and Chlide at 77K were obtained directly from etiolated seedlings in liquid nitrogen under 440-nm excitation with a spectrofluorometer (RF-5300PC; Shimadzu) (Fujii et al., 2017).

### Accession numbers

Sequence data of the genes investigated in this article can be found in The Arabidopsis Information Resource under the following accession numbers: *ACT8* (AT1G49240), *UBQ11* (AT4G05050), *CHLH* (AT5G13630), *CHLD* (AT1G08520), *CHLI1* (AT4G18480), *GUN4* (AT3G59400).

## AUTHOR CONTRIBUTIONS

Akiko Yoshihara performed most experiments, analyzed and interpreted data, and prepared the first draft of the manuscript; Keiko Kobayashi and Noriko Nagata performed transmission electron microscopic experiments, Sho Fujii supported fluoroscopic experiments and edited the drafts of the manuscript, Hajime Wada provided intellectual support and edited the final draft of the manuscript; Koichi Kobayashi designed and conducted experiments, analyzed and interpreted data, and wrote all draft of the manuscript.

## ACKNOWLEDGEMENTS

We thank Tatsuru Masuda (Graduate School of Arts and Sciences, The University of Tokyo) for providing antibody to LPORs.

## SUPPORTING INFORMATION

**Table S1** Oligonucleotide primers used for reverse transcription-quantitative PCR analysis.

**Figure S1** Proportion of total anionic lipids (SQDG + PG) and total plastidic lipids (MGDG + DGDG + SQDG + PG) in total membrane glycerolipids in 4-d-old etiolated seedlings of wild type and anionic lipid mutants.

**Figure S2** Fatty acid compositions of MGDG, DGDG, SQDG, PG, and other phospholipids from 4-d-old etiolated seedlings of wild type and anionic lipid mutants.

**Figure S3** Ultrastructures of etioplasts in *sqd2-2* and *sqd2-2 pgp1-1*.

**Figure S4** Pchlide metabolism in *sqd2-2* and *sqd2-2 pgp1-1*.

**Figure S5.** Quantitative reverse transcription-PCR analysis of the mRNA expression of genes involved in the enzymatic step of Mg insertion into protoporphyrin IX.

**Figure S6** In situ 77K fluorescence spectra in 4-d-old etiolated seedlings of the wild type, *sqd2-2* and *sqd2-2 pgp1-1*.

## REFERENCES

Aoki M, Sato N, Meguro A, Tsuzuki M (2004) Differing involvement of sulfoquinovosyl diacylglycerol in photosystem II in two species of unicellular cyanobacteria. Eur J Biochem 271: 685–693

Babiychuk E, Müller F, Eubel H, Braun H-P, Frentzen M, Kushnir S (2003) Arabidopsis phosphatidylglycerophosphate synthase 1 is essential for chloroplast differentiation, but is dispensable for mitochondrial function. Plant J 33: 899–909

Bligh EG, Dyer WJ (1959) A rapid method of total lipid extraction and purification. Can J Biochem Physiol 37: 911–917

Endo K, Kobayashi K, Wada H (2016) Sulfoquinovosyldiacylglycerol has an essential role in *Thermosynechococcus elongatus* BP-1 under phosphate-deficient conditions. Plant Cell Physiol 57: 2461–2471

Floris D, Kühlbrandt W (2021) Molecular landscape of etioplast inner membranes in higher plants. Nat Plants 7: 514–523

Franck F, Bereza B, Böddi B (1999) Protochlorophyllide-NADP^+^ and protochlorophyllide-NADPH complexes and their regeneration after flash illumination in leaves and etioplast membranes of dark-grown wheat. Photosynth Res 59: 53–61

Franck F, Sperling U, Frick G, Pochert B, van Cleve B, Apel K, Armstrong GA (2000) Regulation of etioplast pigment-protein complexes, inner membrane architecture, and protochlorophyllide a chemical heterogeneity by light-dependent NADPH:protochlorophyllide oxidoreductases A and B1. Plant Physiol 124: 1678–1696

Fujii S, Kobayashi K, Nagata N, Masuda T, Wada H (2017) Monogalactosyldiacylglycerol facilitates synthesis of photoactive protochlorophyllide in etioplasts. Plant Physiol 174: 2183–2198

Fujii S, Kobayashi K, Nagata N, Masuda T, Wada H (2018) Digalactosyldiacylglycerol is essential for organization of the membrane structure in etioplasts. Plant Physiol 177: 1487–1497

Fujii S, Wada H, Kobayashi K (2019) Role of galactolipids in plastid differentiation before and after light exposure. Plants 8: 357

Gabruk M, Mysliwa-Kurdziel B, Kruk J (2017) MGDG, PG and SQDG regulate the activity of light-dependent protochlorophyllide oxidoreductase. Biochem J 474: 1307–1320

Güler S, Seeliger A, Härtel H, Renger G, Benning C (1996) A null mutant of *Synechococcus* sp. PCC7942 deficient in the sulfolipid sulfoquinovosyl diacylglycerol. J Biol Chem 271: 7501–7507

Hagio M, Sakurai I, Sato S, Kato T, Tabata S, Wada H (2002) Phosphatidylglycerol is essential for the development of thylakoid membranes in *Arabidopsis thaliana*. Plant Cell Physiol 43: 1456–1464

Jouhet J (2013) Importance of the hexagonal lipid phase in biological membrane organization. Front. Plant Sci. 4: 494

Kobayashi K, Endo K, Wada H (2016) Roles of lipids in photosynthesis. In Y Nakamura, Y Li-Beisson, eds, Lipids Plant Algae Dev. Subcell. Biochem. Springer International Publishing, Cham, pp 21–49

Kopečná J, Pilný J, Krynická V, Tomčala A, Kis M, Gombos Z, Komenda J, Sobotka R (2015) Lack of phosphatidylglycerol inhibits chlorophyll biosynthesis at multiple sites and limits chlorophyllide reutilization in *Synechocystis* sp. strain PCC 6803. Plant Physiol 169: 1307–1317

Kowalewska Ł, Mazur R, Suski S, Garstka M, Mostowska A (2016) Three-dimensional visualization of the tubular-lamellar transformation of the internal plastid membrane network during runner bean chloroplast biogenesis. Plant Cell 28: 875–891

Li-Beisson Y, Shorrosh B, Beisson F, Andersson MX, Arondel V, Bates PD, Baud S, Bird D, DeBono A, Durrett TP, et al (2013) Acyl-lipid metabolism. Arab Book Am Soc Plant Biol 11: e0161

Masuda T (2008) Recent overview of the Mg branch of the tetrapyrrole biosynthesis leading to chlorophylls. Photosynth Res 96: 121–143

Masuda T, Fusada N, Oosawa N, Takamatsu K, Yamamoto YY, Ohto M, Nakamura K, Goto K, Shibata D, Shirano Y, et al (2003) Functional analysis of isoforms of NADPH:protochlorophyllide oxidoreductase (POR), PORB and PORC, in *Arabidopsis thaliana*. Plant Cell Physiol 44: 963–974

Nguyen HC, Melo AA, Kruk J, Frost A, Gabruk M (2021) Photocatalytic LPOR forms helical lattices that shape membranes for chlorophyll synthesis. Nat Plants 7: 437–444

Okazaki Y, Otsuki H, Narisawa T, Kobayashi M, Sawai S, Kamide Y, Kusano M, Aoki T, Hirai MY, Saito K (2013) A new class of plant lipid is essential for protection against phosphorus depletion. Nat Commun 4: 1510

Okazaki Y, Shimojima M, Sawada Y, Toyooka K, Narisawa T, Mochida K, Tanaka H, Matsuda F, Hirai A, Hirai MY, et al (2009) A chloroplastic UDP-glucose pyrophosphorylase from Arabidopsis is the committed enzyme for the first step of sulfolipid biosynthesis. Plant Cell 21: 892–909

Sato N, Tsuzuki M, Matsuda Y, Ehara T, Osafune T, Kawaguchi A (1995) Isolation and characterization of mutants affected in lipid metabolism of *Chlamydomonas reinhardtii*. Eur J Biochem 230: 987–993

Sato N, Yoshitomi T, Mori-Moriyama N (2020) Characterization and biosynthesis of lipids in *Paulinella micropora* MYN1: Evidence for efficient integration of chromatophores into cellular lipid metabolism. Plant Cell Physiol 61: 869–881

Schindelin J, Arganda-Carreras I, Frise E, Kaynig V, Longair M, Pietzsch T, Preibisch S, Rueden C, Saalfeld S, Schmid B, et al (2012) Fiji: an open-source platform for biological-image analysis. Nat Methods 9: 676–682

Schoefs B (2001) The protochlorophyllide–chlorophyllide cycle. Photosynth Res 70: 257–271

Schoefs B, Franck F (2003) Protochlorophyllide reduction: Mechanisms and evolution. Photochem Photobiol 78: 543–557

Shibata K (1957) Spectroscopic studies on chlorophyll formation in intact leaves. J Biochem (Tokyo) 44: 147–173

Smeller L, Solymosi K, Fidy J, Böddi B (2003) Activation parameters of the blue shift (Shibata shift) subsequent to protochlorophyllide phototransformation. Biochim Biophys Acta 1651: 130–138

Solymosi K, Schoefs B (2010) Etioplast and etio-chloroplast formation under natural conditions: the dark side of chlorophyll biosynthesis in angiosperms. Photosynth Res 105: 143–166

Solymosi K, Smeller L, Ryberg M, Sundqvist C, Fidy J, Böddi B (2007) Molecular rearrangement in POR macrodomains as a reason for the blue shift of chlorophyllide fluorescence observed after phototransformation. Biochim Biophys Acta 1768: 1650– 1658

Sperling U, Franck F, van Cleve B, Frick G, Apel K, Armstrong GA (1998) Etioplast differentiation in Arabidopsis: Both PORA and PORB restore the prolamellar body and photoactive protochlorophyllide–F655 to the cop1 photomorphogenic mutant. Plant Cell 10: 283–296

Tanaka R, Kobayashi K, Masuda T (2011) Tetrapyrrole metabolism in *Arabidopsis thaliana*. Arab Book Am Soc Plant Biol 9: e0145

Tanoue R, Kobayashi M, Katayama K, Nagata N, Wada H (2014) Phosphatidylglycerol biosynthesis is required for the development of embryos and normal membrane structures of chloroplasts and mitochondria in Arabidopsis. FEBS Lett 588: 1680–1685

Wang P, Grimm B (2021) Connecting chlorophyll metabolism with accumulation of the photosynthetic apparatus. Trends Plant Sci 26: 484–495

Wickham H (2009) Elegant graphics for data analysis (ggplot2). Appl. Spat. Data Anal. R

Xu C, Härtel H, Wada H, Hagio M, Yu B, Eakin C, Benning C (2002) The *pgp1* mutant locus of Arabidopsis encodes a phosphatidylglycerolphosphate synthase with impaired activity. Plant Physiol 129: 594–604

Yoshihara A, Kobayashi K (2022) Lipids in photosynthetic protein complexes in the thylakoid membrane of plants, algae, and cyanobacteria. J Exp Bot 73: 2735–2750

Yoshihara A, Nagata N, Wada H, Kobayashi K (2021) Plastid anionic lipids are essential for the development of both photosynthetic and non-photosynthetic organs in *Arabidopsis thaliana*. Int J Mol Sci 22: 4860

Yu B, Benning C (2003) Anionic lipids are required for chloroplast structure and function in Arabidopsis. Plant J 36: 762–770

